# Loss of CHIP Expression Perturbs Glucose Homeostasis and Leads to Type II Diabetes through Defects in Microtubule Polymerization and Glucose Transporter Localization

**DOI:** 10.1101/166389

**Authors:** Holly McDonough, Kaitlin C. Lenhart, Sarah M. Ronnebaum, Chunlian Zhang, Jie An, Andrea Portbury, Christopher B. Newgard, Monte S. Willis, Cam Patterson, Jonathan C. Schisler

## Abstract

Recent evidence has implicated CHIP (carboxyl terminus of Hsc/Hsp70-interacting protein), a co-chaperone and ubiquitin ligase, in the functional support of several metabolism-related proteins, including AMPK and SirT6. In addition to previously reported aging and stress intolerance phenotypes, we find that CHIP ^-/-^ mice also demonstrate a Type II diabetes-like phenotype, including poor glucose tolerance, decreased sensitivity to insulin, and decreased insulin-stimulated glucose uptake in isolated skeletal muscle, characteristic of insulin resistance. In CHIP-deficient cells, glucose stimulation fails to induce translocation of Glut4 to the plasma membrane. This impairment in Glut4 translocation in CHIP-deficient cells is accompanied by decreased tubulin polymerization associated with decreased phosphorylation of stathmin, a microtubule-associated protein required for polymerization-dependent protein trafficking within the cell. Together, these data describe a novel role for CHIP in regulating microtubule polymerization that assists in glucose transporter translocation, promoting whole-body glucose homeostasis and sensitivity to insulin.

## INTRODUCTION

CHIP (carboxyl terminus of Hsc/Hsp70-interacting protein) is a dual function protein, serving both co-chaperone and ubiquitin ligase activities (4, 11-13, 24, 34, 38, 41, 42, 46). CHIP plays an integral role in the maintenance of protein homeostasis largely by regulating protein quality control pathways. CHIP has also been implicated in the aging process, with CHIP-deficient mice demonstrating an accelerated aging phenotype that includes reduced life span, decreased dermal thickness, increased osteoporosis and kyphosis, accumulation of oxidized lipids and misfolded proteins, and early cellular senescence (35). In addition to these hallmarks of increased aging, CHIP-deficient mice are significantly smaller than their wild-type littermates, with reduced whole-body fat stores, decreased subcutaneous adipose layers, and evidence of muscle wasting. These deficits prompted us to examine whether the lack of CHIP protein in these mice may result in dysregulation of metabolic homeostasis, in particular glucose/insulin signaling.

Glucose transport from the blood into fat and skeletal muscle cells is achieved by a carrier-mediated system. The vast majority of glucose uptake occurs in skeletal muscle (27) and is facilitated by a family of protein transporters, including the insulin-sensitive glucose transporter 4 (Glut4). In resting cells, the majority of Glut4 resides in the membrane of intracellular vesicles. However, in response to stimulation by insulin, Glut4 translocates to the cell plasma membrane, where it is exocytosed, allowing glucose to enter the cell by facilitated diffusion. Insulin stimulation greatly enhances the recruitment of Glut4 to the plasma membrane. However, under basal conditions, Glut4 is constantly cycling between the cytoplasm and the cell membrane, and this movement is facilitated by a complex network of cytoskeletal microfilament and microtubule components. Any interference with the integrity of this network results in a failure of Glut4 to move within the intracellular compartment and an inability for the cell to take up glucose.

Given our observation that CHIP-deficient mice develop signs of metabolic dysregulation (in the form of decreased body fat and muscle wasting), along with the fact that many CHIP substrates are involved in the insulin signaling pathway (for example, AKT (15), PTEN (2), and SGK1 (7)), we investigated whether insulin/glucose metabolism was disrupted in CHIP-deficient mice. Indeed, CHIP-deficient mice exhibit numerous signs of dysregulation in these pathways – hyperglycemia, reduced glucose clearance, and reduced glucose uptake in muscle – indicative of insulin resistance. We observe impairments in Glut4 translocation and microtubule polymerization in CHIP-deficient cells. Additionally, CHIP-deficient isolated gastrocnemius myofibers demonstrated inadequate sub-sarcolemmal polymerization of microtubules after a glucose bolus. Together, our results demonstrate that CHIP is involved in the cellular response to both glucose and insulin by contributing to the dynamic response of the microtubule network required for Glut4 translocation, thereby promoting the cellular response to insulin and regulating glucose homeostasis.

## MATERIALS AND METHODS

*Animal care*. Wild-type and CHIP-deficient mice were generated and maintained on a mixed genetic background of C57BL/6 and 129SvEv (C57BL/6 ×129), as previously described (13), or by repeatedly backcrossing the 129SvEv mouse strain (Charles River, Wilmington, MA) with 129Sv/C57B6 mice carrying a single mutated CHIP allele (46, 53). Mice in the present study were 3-3.5 months old. All animal husbandry and experiments were approved by the Institutional Care and Use Committee for Animal Research at The University of North Carolina at Chapel Hill.

*Serum glucose and insulin*. Serum samples were obtained by mandibular punch and collected from “fed” animals (*ad libidum* night feeding) and from “fasted” animals (no food, 16-20 h overnight). Serum glucose was measured with the Glucose Assay Kit (Cayman Chemical Company, Ann Arbor, MI), and serum insulin was measured with the Rat Insulin Enzyme Immunoassay Kit (SPI-BIO Bertin Pharma, Montigny le Bretonneux, France).

*Metabolic studies*. For glucose and insulin tolerance testing, fasted mice were given an intraperitoneal (IP) injection of 20% D-glucose (2 µg glucose/g body mass) or insulin (0.75 U/kg body mass). Blood glucose was measured from tail blood before glucose injection (t = 0) and at the indicated time points after injection. Glucose was measured with a Precision Xtra blood glucose meter (Abbott Diabetes Care Inc., Alameda, CA). Hyperinsulinemic-euglycemic clamp studies were performed on adult mice as previously described (3).

*2-deoxyglucose uptake in skeletal muscle*. Fasted mice received an IP injection of 20% D-glucose (2 µg glucose/g body mass) containing 1 µCi/ml [^3^H] 2-deoxy-D-glucose (PerkinElmer, Waltham, MA). Mice were sacrificed by cervical dislocation 40 min after injection, and muscles were dissected and flash-frozen in liquid nitrogen. Frozen tissue was pulverized and solubilized in 0.1 N NaOH containing 0.1% SDS. 400 µl aliquots of the solubilized muscle were added to 4 ml Ecoscint H scintillation fluid (National Diagnostics, Charlotte, NC) and counted on an LS 6500 multipurpose scintillation counter (Beckman Coulter, Indianapolis, IN); counts were corrected for protein concentration using Protein Assay Dye (Bio-Rad, Hercules, CA) and normalized to wild-type gastrocnemius values per strain of mice.

*Cell culture*. The mouse skeletal muscle-derived C2C12 cell line was transduced with lentiviral particles expressing short hairpin RNAs (shRNA) targeting either CHIP mRNA (shCHIP) or a non-target (shCONT). Briefly, C2C12 cells were infected with lentivirus expressing CHIP shRNA (Open Biosystems, Pittsburgh, PA; catalog numbers TRCN0000007528, TRCN0000008527, and TRCN0000008530; empty vector control pLK0.1 catalog number RHS4080). After 24 h, cells were selected with 2 µg/ml puromycin (Sigma-Aldrich, St. Louis, MO) for one week, a time point at which all uninfected cells had died. The surviving cells were expanded and stored in liquid nitrogen with freezing media. For experiments, cells were cultured in DMEM (Life Technologies, Grand Island, NY) containing 25 mM glucose and 10% FBS (Sigma-Aldrich). Glucose stimulation was performed by incubating cells in serum-free DMEM/5.5 mM glucose for 4 h (indicated by t = 0), then replacing the media with 25 mM glucose DMEM/10% serum/100 nM insulin (Lilly, Indianapolis, IN), and cells were harvested at the indicated time points.

*Immunoblotting, immunocytochemistry, and antibodies*. Cell lysates were collected in RIPA buffer containing 150 mM NaCl, 50 mM Tris HCl pH 7.5, 0.25% deoxycholic acid, 1% NP-40, and 1X protease inhibitor cocktail (Sigma-Aldrich) and subsequently separated into NP-40-soluble and -insoluble fractions by centrifugation. Protein lysates were separated using Bis-Tris 4-12% SDS-PAGE (Life Technologies), transferred to polyvinylidene fluoride membranes (EMD Millipore, Billerica, MA) and immunoblotted using standard chemiluminesence technique. Primary antibodies included: α-tubulin, β-actin, and β-tubulin (Sigma-Aldrich); GAPDH and Glut4 (EMD Millipore); BiP, CHIP, stathmin, and phosphorylated stathmin at serine 16 (Cell Signaling, Danvers, MA); and myc (Santa Cruz Biotechnology, Dallas, TX). For immunocytochemistry, cells were grown on glass slides, washed with PBS, and fixed in 4% paraformaldehyde in PBS (w/v). GFP was visualized directly to detect myc-Glut4-GFP. Indirect immunofluorescence of endogenous Glut4 was carried out by incubating the Glut4 antibody (1:200) and visualizing with an AlexaFluor488 secondary antibody (Life Technologies). In some instances, cells were stained with TexasRed-X Phalloidin (Life Technologies) for 45 min. For indirect immunofluorescence using *ex vivo* muscle fibers, the fixed fibers were transferred to blocking buffer containing 50 mM glycine, 0.25% BSA, 0.04% TritonX-100, and 0.05% sodium azide in PBS, and permeabilized in buffer containing 1% BSA and 0.5% TritonX-100 in PBS for 30 min, after which time fibers were incubated overnight with the α-tubulin antibody diluted 1:1000 in blocking buffer. After 3 PBS washes of 30 min each, the myofibers were incubated for 2 h with an AlexaFluor 488-conjugated goat anti-mouse antibody (Life Technologies) and washed 3 times in PBS for 5 min each. All cells and muscle fiber preparations were mounted with glass cover slips using Vectashield mounting media with DAPI (Fisher Scientific, Pittsburgh, PA). Micrographs were obtained on a Nikon Eclipse E800 upright fluorescent microscope utilizing QCapture software (QImaging Corp., Surrey, British Columbia, Canada).

*OPD assay*. O-phenylenediamine dihydrochloride (OPD) assays were performed using SIGMAFAST OPD tablets according to manufacturer’s instructions (Sigma-Aldrich). Briefly, C2C12 cells in 6 well plates were washed with PBS, fixed for 3 min in 4% paraformaldehyde (w/v) at room temperature, then neutralized with 1% glycine (w/v) in PBS at 4°C for 10 min. Cells were blocked for 30 min at 4°C with PBS containing 10% goat serum and 3% BSA, incubated with anti-myc antibody at 4°C for 30 min, then incubated with peroxidase-conjugated goat anti-mouse secondary antibody. Following removal of the secondary antibody, 1 ml of O-phenylenediamine dihydrochloride in phosphate-citrate buffer with urea hydrogen peroxide (Sigma-Aldrich) was added to each well of cells for 20 min. The reaction was stopped by adding 0.25 ml of 3 M HCl. Optical absorbance of the supernatant was measured at 492 nm on a SmartSpec 3000 (Bio-Rad). The myc epitope on the exofacial loop of the membrane-inserted myc-Glut4-GFP is detected first by the anti-myc antibody, then by the secondary horseradish peroxidase-conjugated antibody. The action of the peroxidase secondary antibody on OPD changes its absorbance. The change in absorbance correlates with the amount of exposed myc epitope and thus can be used to calculate the amount of membrane-inserted myc-Glut4-GFP.

*Two-dimensional differential in gel electrophoresis (2D-DIGE), matrix-assisted laser desorption ionization time of flight (MALDI-TOF) mass spectrometry, and functional clustering*. Differential protein expression using 2D-DIGE comparing wild-type and CHIP-deficient mouse embryonic fibroblast protein extracts was carried out as previously described (44). 34 spots identified via 2D-DIGE (unpaired t-test, *p* < 0.05, FDR < 25%) were identified via MALDI-TOF mass spectrometry and analyzed using the DAVID functional annotation clustering tool (14, 21). The mean fold-enrichment (log_2_) of each functional cluster (x-axis) and corresponding proteins (y- axis) were analyzed using a Pearson’s centered, complete linkage clustering analysis as previously described (8).

*Single muscle fiber preparation*. Wild-type and CHIP-deficient mice were fasted overnight and split into two groups. The first group of mice maintained the fast, whereas the second received an IP injection of 20% D-glucose (2 µg glucose/g body mass). After 40 min, mice were sacrificed by cervical dislocation, and the gastrocnemius was immediately removed and placed in 2% paraformaldehyde (w/v) for 1 h, then rinsed several times in PBS. Small bundles of 1-3 fibers were then teased away with fine forceps.

*Statistics*. Statistical tests were performed as indicated in the methods, figure legends, and table legends.

## RESULTS

### CHIP is necessary for optimal glucose homeostasis

CHIP-deficient mice demonstrate an accelerated aging phenotype as well as other maladies, including muscle wasting, that suggest metabolic imbalances (35). Given the high level of CHIP expression detected in human skeletal muscle (4), the association between muscle atrophy and altered glucose homeostasis (47), and the established link between the loss of muscle mass in the elderly with Type II diabetes (5, 16), we hypothesized that CHIP-deficient mice may have altered glucose homeostasis stemming from skeletal muscle dysfunction. Consistent with our hypothesis, mice lacking CHIP exhibited a mild hyperglycemia that was more pronounced under fasted conditions (Fig. 1*A*). The change in glucose homeostasis was accompanied by a trend towards an increase in insulin levels that was unmasked in the fed state (Fig. 1*A*), suggesting that decreased insulin responsiveness in the peripheral tissues of CHIP-deficient mice may result in compensatory hyperinsulinemia typical of Type II diabetes (18). To test this, a glucose tolerance test was performed on wild-type and CHIP-deficient 129SvEv mice to determine the acute ability of tissues in these mice to remove glucose from the blood. CHIP-deficient mice failed to clear glucose from the blood efficiently compared to wild-type mice (48 ± 9% increase in the AUC comparing CHIP ^-/-^ to wild-type mice, Fig. 1*B*). To measure whether the defect in glucose tolerance was due to a reduction in insulin release or insulin receptor sensitivity, an insulin tolerance test was performed. We found that intraperitoneal administration of insulin had a smaller effect on lowering blood glucose levels in CHIP-deficient mice at all time points tested (49 ± 6% increase in the AUC comparing CHIP ^-/-^ to wild-type mice, Fig. 1*C*), suggesting that insulin sensitivity is impaired in CHIP ^-/-^ mice. To directly assess insulin sensitivity, we performed a hyperinsulinemic-euglycemic clamp assay. During insulin infusion, CHIP-deficient mice required half the rate of glucose infusion compared to wild-type mice to maintain blood glucose levels (Table 1), indicating whole-body insulin resistance. This finding was supported by the fact that CHIP-deficient mice were unable to suppress hepatic glucose production to the same extent as wild-type mice, and there was a dramatic reduction in the ability of CHIP-deficient gastrocnemius to take up 2-deoxyglucose during hyperinsulimia (45 ± 9%; Table 1). This poor glucose tolerance phenotype and impaired glucose uptake in skeletal muscle was recapitulated in CHIP ^-/-^ mice bred on a mixed C57BL/6 × 129 background (Fig. 1*D* and Table 1). Taken together, these data demonstrate that the absence of CHIP leads to a Type II diabetes-like phenotype under physiological conditions due to impaired glucose uptake and insulin resistance in skeletal muscle.

**Table 1.** Results of hyperinsulinemic-euglycemic clamp study and glucose uptake in muscle.

| **129SvEv** | **Wild-type** | **CHIP<sup>-/-</sup>** | ***p*** |
| --- | --- | --- | --- |
| Glucose infusion rate (mg/kg/min) | 22 ± 4 | 11 ± 1 | 0.0018 |
| Hepatic glucose output (mg/kg/min) | 13 ± 3 | 21 ± 4 | 0.0186 |
| Insulin-induced suppression of hepatic glucose production (%) | 78 ± 6 | 41 ± 10 | 0.0007 |
| Gastrocnemius 2-deoxyglucose uptake (relative counts/mg) | 100 ± 8 | 55 ± 9 | 0.0022 |
| **mixed C57BL/6 × 129SvEv** | **Wild-type** | **CHIP<sup>-/-</sup>** | ***p*** |
| Gastrocnemius 2-deoxyglucose uptake (relative counts/mg) | 100 ± 25 | 21 ± 4 | 0.0137 |
| Soleus 2-deoxyglucose uptake (relative counts/mg) | 916 ± 161 | 393 ± 47 | 0.0140 |

**Fig. 1.**
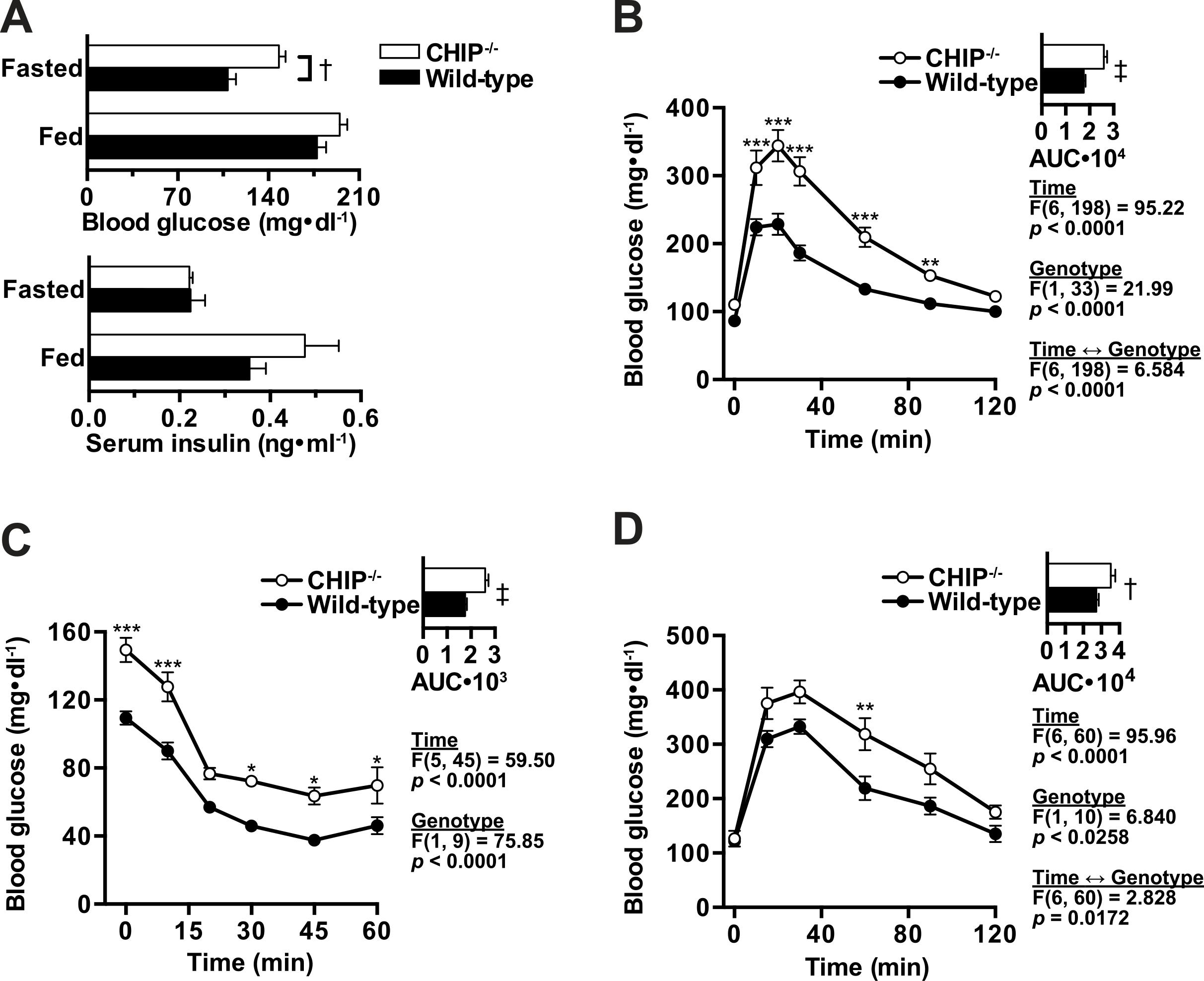
Mice deficient in CHIP expression demonstrate Type II diabetic phenotypes. *A*: *ad libitum* fed or overnight-fasted blood glucose (upper) and insulin (lower) in 129SvEv mice represented by the mean ± SEM (n > 8 per genotype, per condition): † *p* = 0.00018 via unpaired t-test comparing wild-type and CHIP ^-/-^ mice. *B*-*C*: blood glucose levels in 129SvEv mice measured at the indicated time points after (*B*) glucose or (*C*) insulin bolus injection represented by the mean ± SEM (n = 15/20 or 5/6 in wild-type/CHIP ^-/-^ mice for glucose or insulin tolerance, respectively). *D*, 2-deoxyglucose uptake in gastronemius of 129SvEv mice (relative counts/mg).

### CHIP is necessary for optimal Glut4 translocation to the membrane following insulin stimulation

We hypothesized that the severe defect in glucose uptake associated with CHIP deficiency (Fig. 1*B-D*, Table 1) may be linked to the inability of Glut4, the major glucose transporter found in skeletal muscle and adipose tissue, to translocate from the cell interior to the plasma membrane in response to insulin stimulation, thereby hindering the cellular uptake of glucose (22). Following insulin stimulation, a series of intracellular signaling pathways promote Glut4 movement from intracellular vesicles to the cell membrane (9). Once Glut4 reaches the cell membrane, it fuses and undergoes exocytosis to facilitate the diffusion of glucose through the cell membrane. Glut4 translocation is essential for successful insulin signaling and glucose uptake by skeletal muscle cells (23, 26, 39, 45, 51). We employed lentiviral transduction of control or CHIP shRNA in mouse skeletal muscle C2C12 cells (shCONT and shCHIP cell lines) to investigate if a reduction in CHIP expression influences insulin-dependent Glut4 translocation. To measure Glut4 translocation, we transfected shCONT and shCHIP C2C12 cells with a myc-Glut4-GFP construct (myc N-terminus, GFP C-terminus; (52)) and examined the location of Glut4-GFP protein following 4 h serum starvation before and after a challenge with 25 mM glucose and 100 nM insulin. After serum starvation and prior to glucose/insulin stimulation (time 0), Glut4-GFP was largely localized to the perinuclear region in both the shCONT and shCHIP C2C12 cells (Fig. 2*A*). However, following 30 min of insulin stimulation, Glut4-GFP translocated to the membrane in shCONT cells, but remained in a perinuclear region in the shCHIP cells (Fig. 2*A*). To verify this lack of Glut4 movement in response to insulin in shCHIP cells, we took advantage of the myc tag on the Glut4-GFP construct and performed a colorimetric assay of surface myc-Glut4-GFP using O-phenylenediamine dihydrochloride (OPD). In this method, insertion of myc-Glut4-GFP protein into the cell membrane results in the exofacial positioning of the myc tag. This tag is subsequently labeled with a secondary antibody conjugated to peroxidase and then is treated with OPD reagent, which results in a colorimetric reaction that can be monitored by measuring light absorbance at 492 nm (50). Using this assay, we confirmed that, whereas the membrane level of myc-Glut4-GFP increased following insulin stimulation in shCONT cells, the amount of membrane-bound myc-Glut4-GFP did not change following insulin stimulation in shCHIP cells (Fig. 2*B*). This lack of Glut4 translocation to the membrane in the shCHIP cells suggests that CHIP is involved in insulin-dependent trafficking of Glut4 to the cell membrane.

**Fig. 2.**
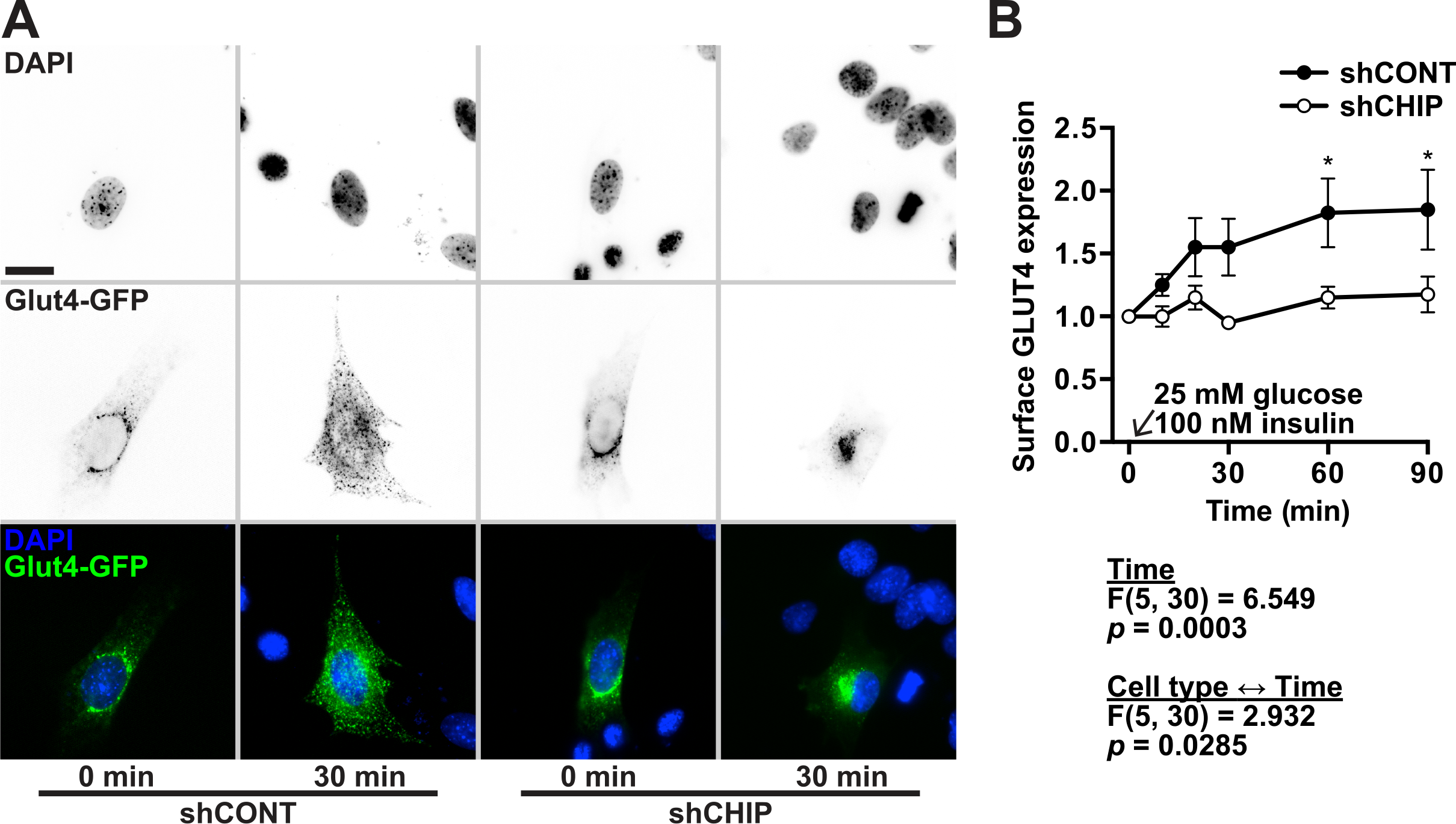
C57BL/6x129 mice (mixed background) also display Type II diabetic phenotype. *A:* overnight fasted mice blood glucose levels measured at the indicated time points after a glucose bolus injection represented by the mean ±SEM (n=4males, 2females per genotype)*B*. Glucose stimulated insulin secretion in freshly isolated islets from wild-type and CHIP ^-/-^ mice. *C*. 2-deoxyglucose uptake in gastronemius and *D*. in soleus of overnight fasted C57BL/6x129 mice forty minutes after IP glucose bolus.

### CHIP deficiency results in cytoskeletal abnormalities in shCHIP C2C12 cells

The movement of Glut4 to the cell membrane requires the coordinated assembly of both actin microfilaments and tubulin-containing microtubules (17, 36, 37, 43). Therefore, to determine the possible cause for aberrant Glut4 translocation in CHIP-deficient cells, we first examined whether the arrangement and insulin-mediated response of filamentous actin (F-actin) was altered in CHIP-deficient C2C12 cells. In unstimulated cells, F-actin filaments were evenly distributed throughout both shCONT and shCHIP cells, running parallel to the longitudinal axis of the cell (Fig. 3*A*). Following insulin stimulation for 30 min, F-actin foci localized near the plasma membrane edge in both shCONT and shCHIP cells (Fig. 3*A*). In the shCONT cells, the F-actin foci co-stained for Glut4, supporting previous reports that actin is critical for the successful fusion and exocytosis of Glut4 at the plasma membrane (32). In contrast, although some plasma membrane-localized F-actin foci were present in shCHIP cells stimulated with insulin, Glut4 did not colocalize with these structures. Instead, we saw a heavy concentration of F-actin/Glut4 foci clustered around the nucleus in insulin-stimulated shCHIP cells. (Fig. 3*A*). The patterning of these structures was reminiscent of the fractured Golgi associated with microtubule depolymerization in neurons (19), suggesting that the microtubule network responsible for ensuring the successful transport of Glut4 to the plasma membrane in response to insulin stimulation may be compromised in shCHIP cells. Remodeling of the microtubule network within cells is critical for Glut4 dependent insulin-induced glucose uptake (10, 17, 31). The signaling pathways downstream of insulin stimulation result in the polymerization of soluble tubulin dimers, leading to the formation of insoluble microtubules. With this in mind, we took advantage of the solubility dynamic of the microtubule protein β-tubulin to investigate whether the microtubule network formed in response to insulin stimulation is disrupted in CHIP-deficient cells. After 10 min of insulin stimulation, tubulin began appearing in the NP-40-insoluble pellet fraction in lysates collected from shCONT cells (Fig. 3*B*-*C*). In contrast, no movement of tubulin to the insoluble fraction was detected in shCHIP cells, even at later time points. Importantly, these data indicate a severe disruption in the ability of tubulin to polymerize in response to insulin in CHIP-deficient C2C12 cells.

**Fig. 3.**
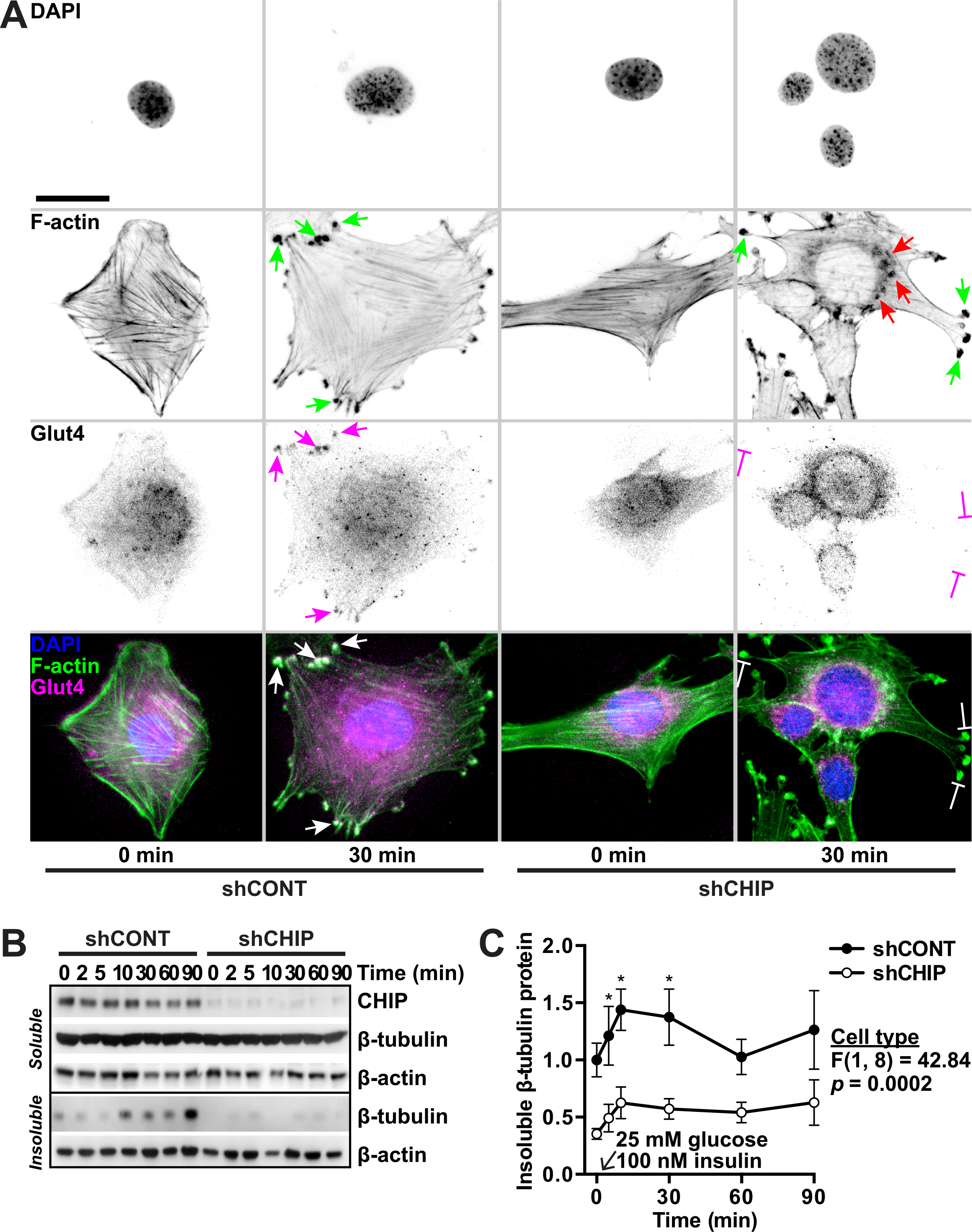
Diagram of “components” of glucose metabolism showing roles of liver and skeletal muscle.

### Stathmin phosphorylation is decreased in CHIP-deficient C2C12 cells

Because our data suggested that tubulin polymerization is impaired in the absence of CHIP, we investigated whether other cytoskeletal-associated proteins were reliant on CHIP expression. First, we identified proteins whose steady-state protein levels were affected by CHIP expression by performing two-dimensional difference gel electrophoresis (2D-DIGE) on protein extracts from primary wild-type and CHIP-deficient MEFs. We identified *Actin Cytoskeleton and Morphogenesis* as one of the major classes of proteins differentially regulated by CHIP expression (Fig. 4*A*). Included in this group of proteins was stathmin, a microtubule-associated protein that regulates the polymerization of tubulin (48). Stathmin levels were reduced in the shCONT cells compared to the CHIP-deficient cells, suggesting that CHIP may directly affect stathmin cellular function. Stathmin activity is regulated by both total levels and by phosphorylation. Unphosphorylated stathmin binds to the α- and β-tubulin dimers and helps to maintain a pool of tubulin readily available for polymerization in the soluble fraction, but phosphorylation of stathmin prevents stathmin from binding to tubulin dimers, thereby allowing tubulin polymerization (33, 40, 48). To determine if the phosphorylation of stathmin is impacted by CHIP expression in a cellular context where tubulin reorganization is triggered, we measured stathmin serine 16 phosphorylation after serum starvation and subsequent insulin stimulation (33). Following insulin stimulation, the level of phosphorylated stathmin was increased in shCONT cells (Fig. 4*C*-*D*), coincident with the appearance of polymerized tubulin (Fig. 3*B*-*C*). In contrast, a much smaller increase in phosphorylated stathmin was seen in shCHIP cells, in agreement with an observed lack of polymerized tubulin in these cells. Taken together, the decreased insulin sensitivity in CHIP-deficient mice and insufficient Glut4 translocation may be caused by impaired stathmin phosphorylation and inadequate tubulin polymerization.

**Fig. 4.**
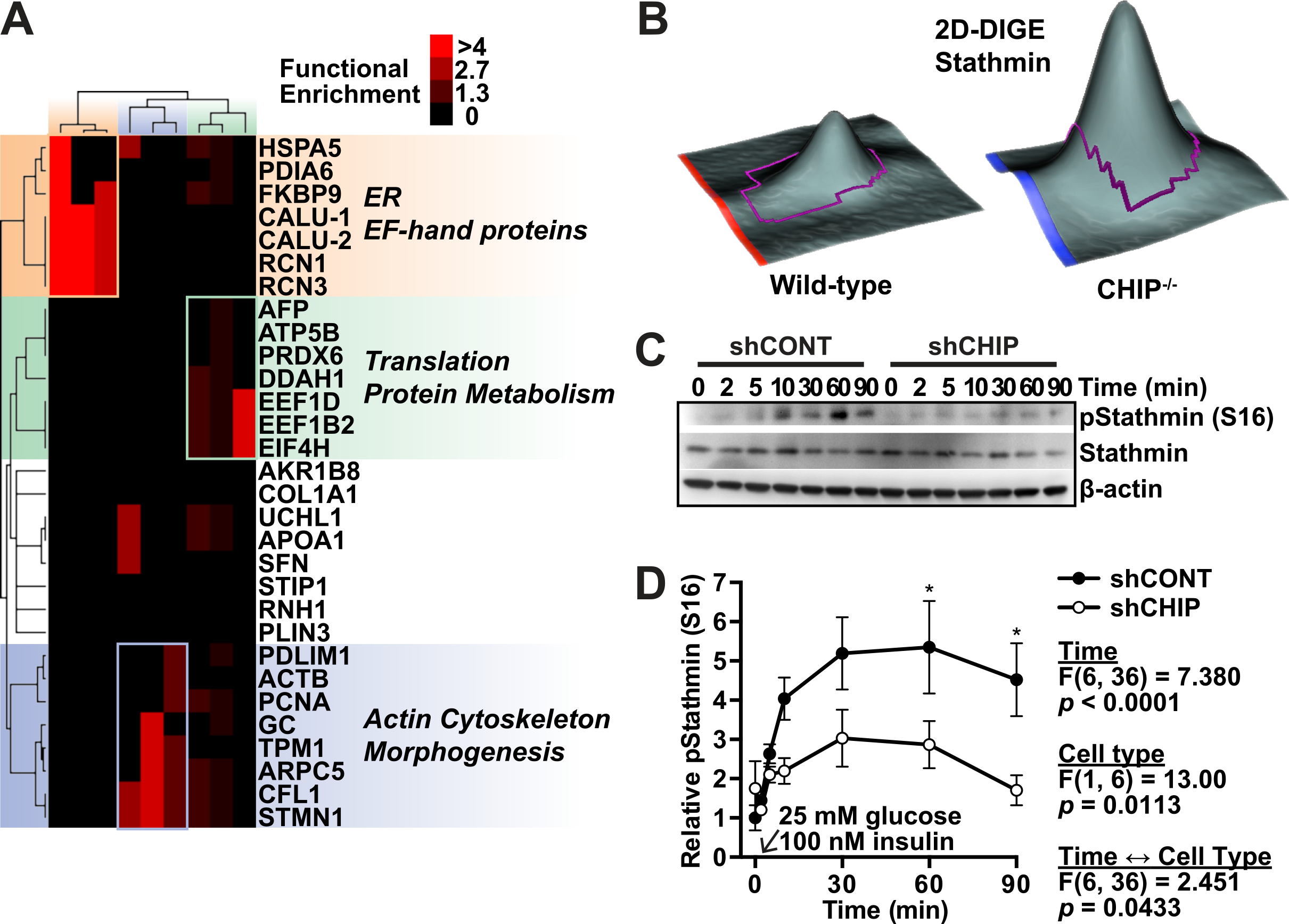
Absence of CHIP in C2C12 cells affects insulin stimulated glucose metabolism. *A*. glucose uptake in control (shCONT) and CHIP deficient (shCHIP) C2C12 cells, measured at the indicated times after 4 hours serum starve, then complete media plus insulin (shCONT−,shCHIP--). *B*. glucose uptake in CHIP deficient (shCHIP) C2C12 cells infected with AdGFP(--) or AdCHIP(−), measured at the indicated times. *C*. microarray data from wild-type and CHIP deficient primary MEFs. Hierarchical functional clustering of 34 differentially expressed proteins identified via 2D-DIGE analysis comparing wild-type and CHIP ^-/-^ fibroblast protein extracts. Proteins (rows) are represented by their functional classification in the indicated clusters (columns): orange, ER and EF-hand proteins; green, translation and protein metabolism; and blue, actin cytoskeleton and morphogenesis. The heat map indicates the fold enrichment of each cluster compared to the relative enrichment that occurs in the mouse proteome.

### CHIP is necessary for optimal microtubule polymerization *in vivo* following insulin stimulation

To determine if the lack of microtubule polymerization in CHIP-deficient C2C12 cells was recapitulated in an *in vivo* setting, we examined microtubule polymerization in freshly isolated myofibers from CHIP-deficient mice. In response to insulin stimulation, microtubules within individual muscle fibers longitudinally span myofibrils just beneath the sarcolemma and transversely across the myofibrils, with a concentration in the area of the I-band leading to a lattice-like microtubule appearance (25). Microtubules are also found in the perinuclear region and radiate from the perinuclear region to run transversely into the interior of the myofibril, an arrangement that is purported to support intracellular transport (25). To examine whether the microtubule arrangement is aberrant in CHIP-deficient muscle fibers following insulin stimulation, we fasted wild-type and CHIP-deficient mice overnight and then challenged both genotypes with an IP bolus of glucose. Neither wild-type nor CHIP-deficient myofibers displayed substantial sub-sarcolemmal nor perinuclear α-tubulin staining after the overnight fast. However, after an injection of glucose, the wild-type fibers exhibited intense perinuclear α-tubulin staining and a striking lattice arrangement of α-tubulin-positive fibers at the surface of the sarcolemma (Fig. 5). In contrast, this lattice appearance was strikingly absent in CHIP-deficient myofibers, revealing a lack of microtubule polymerization at the myofiber surface (Fig. 5). Taken together, we observe hyperglycemia and insulin resistance in CHIP-deficient mice, and we demonstrate that shCHIP cells exhibit decreased microtubule polymerization and Glut4 translocation. We propose that CHIP plays a critical role in promoting insulin-stimulated microtubule polymerization and the movement of Glut4 to the membrane, facilitating the import of glucose into the cell and maintaining glucose homeostasis.

**Fig. 5.**
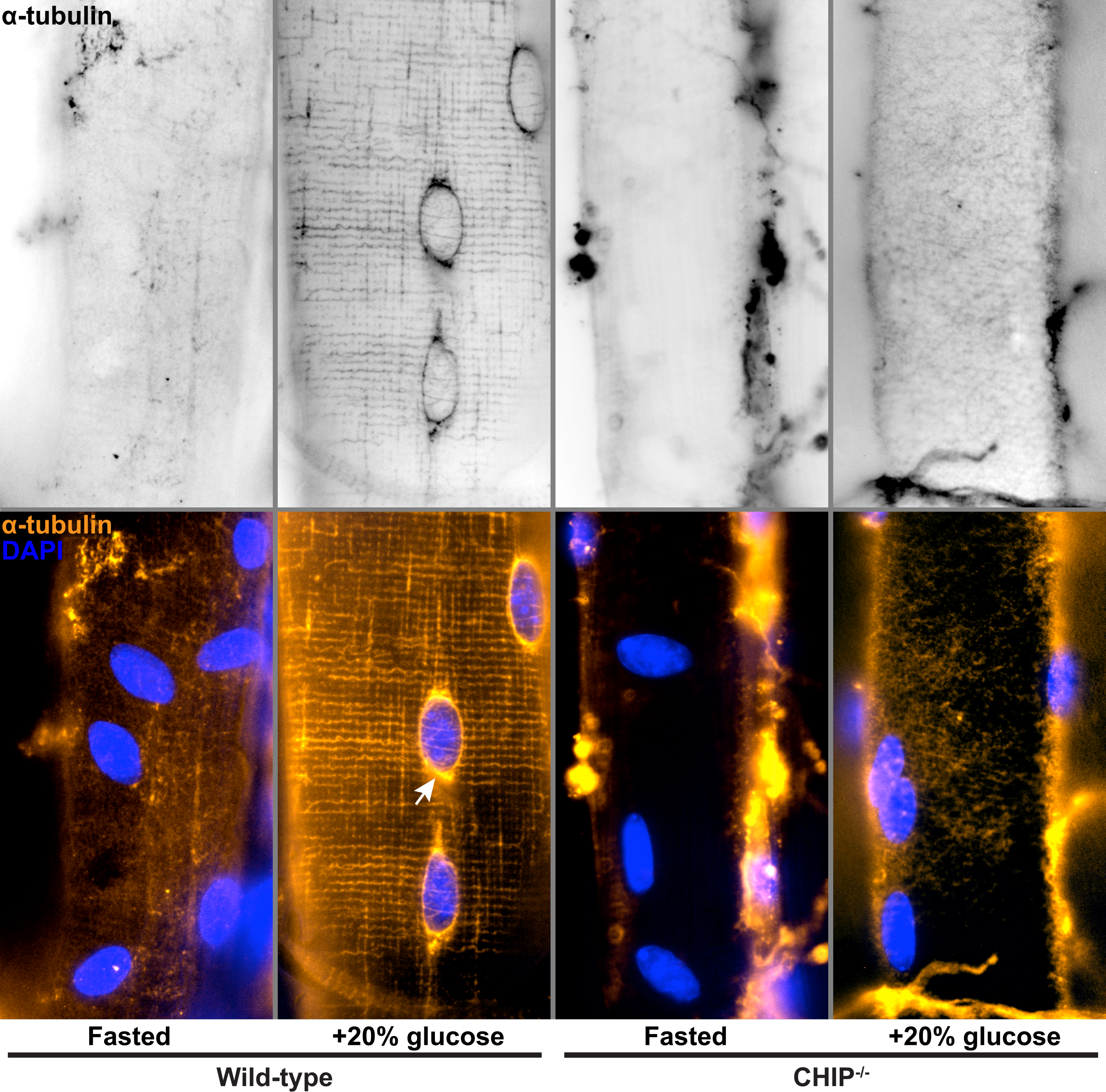
Cytoskeletal defects in cells with reduced CHIP expression. *A*: fluorescent micrographs of C2C12 control cells (shCONT) or cells with reduced CHIP expression (shCHIP). After serum starvation (0 min) or stimulation (30 min) with complete media plus insulin, cells were fixed and visualized for nuclei (DAPI), phalloidin staining (F-actin), and endogenous Glut4 expression. Red arrows indicate perinuclear F-actin foci; green arrows indicate membrane edge F-actin foci; white arrows indicate F-actin/Glut4 colocalization. False color overlays are included. Scale bar indicates 10 µm. *B*. Immunoblot for Bip in C2C12 cells after 4 hrs serum starve, then 30 min in complete media plus insulin.

## DISCUSSION

Glucose transporter translocation is an important cellular mechanism in glucose homeostasis, especially in insulin-sensitive tissues such as skeletal muscle. Previously, we reported that mice lacking CHIP expression exhibit phenotypes such as accelerated aging and decreased tolerance to cardiac stressors such as ischemia reperfusion and pressure overload (35). Here we add to the characterization of the CHIP-deficient phenotype by demonstrating that CHIP-deficient mice also develop a Type II diabetic phenotype, including glucose intolerance (Fig. 1*B*, 1*D*), insulin resistance (Fig. 1*C*, Table 1), and decreased glucose uptake in skeletal muscle (Table 1). The apparent cause for this aberrant glucose/insulin response is at least in part due to the disruption of microtubule reorganization in response to insulin stimulation (Fig. 3*A*, 3*B*, 3*C*, and 5) that is necessary for the glucose transporter Glut4 to translocate to the cell membrane and facilitate the entry of glucose into the cell (Fig 2*A*, 2*B*, and 3*A*). These data support the hypothesis that CHIP expression is necessary for maintaining optimal glucose/insulin signaling and provides further evidence of an intersection between protein quality control and metabolic pathways.

The idea that CHIP, a co-chaperone/ubiquitin ligase involved in protein quality control pathways, could play a role in metabolic homeostasis evolved over the last several years. The discovery that CHIP forms a complex with the stress-induced kinase SGK1 (serum- and glucocorticoid-regulated kinase-1; (6)), resulting in the inhibition of SGK kinase activity, positions CHIP to play an integral role in modifying myriad cellular responses to stress, including insulin signaling (20, 30). CHIP also influences the activity of the peroxisome proliferator-activated receptor (PPAR) proteins PPARα and PPARβ via its interaction with Hsp90 (49). The PPAR proteins are nuclear receptor proteins that regulate the expression of numerous genes involved in cellular differentiation, development, and metabolism (1). By binding to Hsp90, CHIP induces the activation of PPARα that, among other things, impacts PPARα’s regulation of fatty acid catabolism as well as amino acid and carbohydrate metabolism (28, 29). Perhaps the most direct link between CHIP and metabolic regulation is the recent discovery that the energy sensing enzyme AMP-activated kinase (AMPK) is a physiological substrate of CHIP’s autonomous chaperone activity (46). CHIP promotes phosphorylation of AMPK via LKB1, thereby increasing AMPK activity in response to a decrease in ATP; in addition, CHIP alters the tertiary structure of the α catalytic subunit of AMPK, resulting in enhanced stability and activation of AMPK function (46). Given the complex physiology of glucose homeostasis that requires both tissues and organ systems to work individually as well as coordinately, it is likely that multiple CHIP-dependent pathways including the regulation of cytoskeletal architecture contribute to the phenotype observed in CHIP ^-/-^ mice. Future studies that utilize tissue-specific manipulations of CHIP expression will undoubtedly help elucidate the role of CHIP in tissues with vastly different metabolic properties, especially in complex diseases such as diabetes and heart failure.

Our investigation into possible mechanisms by which glucose homeostasis is disrupted in CHIP-deficient led us to discover a microtubule polymerization defect in the CHIP-deficient cells. The data shows that insulin-responsive microtubule polymerization is impaired in CHIP-deficient skeletal muscle cells and in the myofibers of CHIP-deficient mice. Normally, upon insulin stimulation, the microtubule-associated protein stathmin is phosphorylated, causing stathmin disengagement from tubulin dimers in the cytosol and freeing tubulin to polymerize (33, 40, 48). Upon insulin stimulation, the absence of CHIP abrogates the phosphorylation of serine 16 of stathmin, suggesting the possibility of a role for CHIP at this point in the insulin signaling phosphorylation cascade. CHIP could be acting purely as a chaperone to stathmin, such that the interaction causes a conformational change in stathmin that facilitates phosphorylation, similar to CHIP’s chaperone function in the phosphorylation of AMPK (46). Of course, CHIP could also be acting further upstream. There are two candidate CHIP substrates in this regard, AKT (15) and PTEN (2). To promote the insulin-stimulated phosphorylation cascade, CHIP could have a role as a chaperone to AKT in a conformational-changing interaction that promotes AKT phosphorylation. Or, CHIP’s ubiquitin ligase activity might target PTEN, preventing its phosphatase activity on PIP_3_, thereby prolonging PIP_3_ phosphorylation signaling. Interestingly, we have observed a similar abrogation of Glut4 translocation in AICAR-stimulated C2C12 cells (data not shown), suggesting that CHIP’s effect occurs downstream of the point where the AMPK and the insulin signaling pathways converge (9), which would suggest that CHIP acts downstream of AKT and PTEN. Indeed, insulin-stimulated membrane ruffling still occurs in the CHIP-deficient cells, suggesting that plasma membrane-associated molecules are not affected by the absence of CHIP. Alternatively, CHIP could be targeting an as-yet unidentified protein in insulin signaling that plays a role in the transmission of the phosphorylation cascade to the interior of the cell, a process that remains incompletely understood. Nevertheless, the effect on microtubule polymerization in CHIP-deficient muscle cell lines and muscle cells is striking, and we believe this inhibition of microtubule polymerization causes reduced glucose uptake in cells as well as in skeletal muscle *in vivo*.

**Fig. 6.** CHIP expression is necessary for insulin-stimulated Glut4 localization. *A*: fluorescent micrographs of C2C12 control cells (shCONT) or cells with reduced CHIP expression (shCHIP) transfected with myc-Glut4-GFP cDNA. After serum starvation (0 min) or stimulation (30 min) with complete media plus insulin, cells were fixed and visualized for nuclei (DAPI) and Glut4 expression. False color overlays are included. Scale bar indicates 10 µm. *B*: relative levels of surface Glut4 expression measures by O-phenylenediamine dihydrochloride absorbance in shCONT and shCHIP cells at the indicated time points after stimulation are represented by the mean ± SEM (n = 4): significant results of two-way ANOVA (α = 0.05) are indicated, * *p* < 0.05 via Sidak’s multiple comparisons test comparing shCONT and shCHIP cells at the indicated time point.

**Fig. 7.** CHIP-dependent effects on tubulin polymerization *A*. representative immunoblot analysis of CHIP as well as both soluble and insoluble β-tubulin and β-actin. *B*. densitometry of relative insoluble β-tubulin levels in protein extracts from serum-starved C2C12 cells (time = 0) and at the indicated time points (min) after stimulation with complete media plus insulin represented by the mean ± SEM (n = 4): significant results of two-way ANOVA (α = 0.05) are indicated, * *p* < 0.05 via Sidak’s multiple comparisons test comparing shCONT and shCHIP cells at the indicated time point. *C*: representative immunoblot analysis of phosphorylated stathmin at serine 16 (S16), total stathmin, and β-actin, and *D*: densitometry of relative phosphorylated stathmin (S16) levels in protein extracts from serum-starved C2C12 cells (time = 0) and at the indicated time points (min) after stimulation with complete media plus insulin represented by the mean ± SEM (n = 4): significant results of two-way ANOVA (α = 0.05) are indicated, * *p* < 0.05 via Sidak’s multiple comparisons test comparing shCONT and shCHIP cells at the indicated time point.

**Fig. 8.** Reduced CHIP affects insulin pathway signaling in tissue *A*.immunoblot of indicated insulin pathway molecules comparing wild-type and CHIP deficient tissues *B*. immunoblot for AKT and PTEN in insulin stimulated shCONT and shCHIP C2C12 cells.

**Fig. 9.** CHIP PTEN interaction correlated with sustained PIP3 levels in shCONT C2C12 cells. *A*. Immunoblot of PTEN immunoprecipitated lysate from insulin stimulated shCONT C2C12 Cells. *B*. immunoblot of membrane and cytoplasmic fractions of insulin stimulated shCONT and shCHIP C2C12 cells. *C*. comparing PIP3 levels in insulin stimulated shCONT and shCHIP C2C12 cells.

**Fig. 10.** Differentiated C2C12 myotubes exhibit the same tubulin and Glut4 CHIP effects seen in the undifferentiated cells. *A*. shCONT myotubes exhibit increased glucose uptake in insulin stimulated myotubes compared with CHIP-deficient (shCHIP) myotubes (representative experiment). *B*. immunoblot for tubulin in the soluble and pellet fractions of insulin stimulated shCONT and CHIP-deficient (shCHIP) myotubes. *C*. immunoblot for soluble and pellet tubulin in insulin stimulated AdGFP and AdCHIP infected CHIP deficient (shCHIP) myotubes. *D*. 30 min insulin stimulation of shCONT and CHIP-deficient (shCHIP) myotubes immunobloted for endogenous Glut4.

**Fig. 11.** CHIP expression is necessary for glucose-mediated cytoskeletal dynamic in *ex vivo* skeletal muscle. Representative fluorescent micrographs of individual gastrocnemius muscle fibers isolated from wild-type and CHIP-deficient fasted mice or fasted mice given a glucose bolus (40 min) were stained for α-tubulin (top). False color overlays of α-tubulin (orange) and the nuclear counterstain DAPI (blue) are provided (bottom).

## ACKNOWLEDGEMENTS

We are grateful to the Michael Hooker Proteomics Center at UNC for protein identification services. The current affiliation for Cam Patterson is Presbyterian Hospital/Weill-Cornell Medical Center, New York, NY, USA.

## AUTHOR CONTRIBUTIONS

H.M., C.Z., C.B.N., C.P., and J.C.S. conception and design of research; H.M., C.Z., and J.A. performed experiments; H.M., C.Z., J.A., and J.C.S. analyzed data; H.M., K.C.L., C.Z., C.B.N., M.S.W, C.P., and J.C.S. interpreted results of experiments; H.M., K.C.L., and J.C.S. prepared figures; H.M., K.C.L., S.M.R., A.P., and J.C.S. drafted manuscript; H.M., K.C.L., S.M.R., A.P., M.S.W., and J.C.S. edited and revised manuscript; H.M., K.C.L., S.M.R., C.Z., J.A., A.P., C.B.N., M.S.W., C.P., and J.C.S. approved final version of manuscript.

## DISCLOSURES

The authors have no conflicts of interest to disclose.

Data are represented by the mean ± SEM (n = 4 and 5 per genotype, in 129SvEv and mixed mice, respectively): *p* values from unpaired t-test comparing wild-type and CHIP ^-/-^ mice are provided.

